# Boiler: Lossy compression of RNA-seq alignments using coverage vectors

**DOI:** 10.1101/040634

**Authors:** Jacob Pritt, Ben Langmead

## Abstract

We describe Boiler, a new software tool for compressing and querying large collections of RNA-seq alignments. Boiler discards most per-read data, keeping only a genomic coverage vector plus a few empirical distributions summarizing the alignments. Since most per-read data is discarded, storage footprint is often much smaller than that achieved by other compression tools. Despite this, the most relevant per-read data can be recovered; we show that Boiler compression has only a slight negative impact on results given by downstream tools for isoform assembly and quantification. Boiler also allows the user to pose fast and useful queries without decompressing the entire file. Boiler is free open source software available from github.com/jpritt/boiler.

## INTRODUCTION

Sequence Alignment/Map (SAM/BAM) (22) is a ubiquitous file format for storing RNA (and DNA) sequencing read alignments. For each aligned read, SAM/BAM stores the alignment’s location, shape (described by the CIGAR string), base and quality sequences, and other data. BAM files are the binary equivalent of SAM, and BAM files are often sorted along the genome and compressed.

A drawback of SAM/BAM, and of any format that stores data on a per-read basis, is that file size grows close to linearly with the number of reads. But as sequencing continues to improve (27), and as public archives fill with more datasets, the burden of storing aligned sequencing data also increases rapidly. The Sequence Read Archive (20), which stores raw sequencing reads, grew from 3 to 4 petabases from February to August of 2015. It is increasingly common for RNA-seq studies to span hundreds or thousands of samples, with tens of millions of reads per sample (2, 17).

Compressed formats eliminate redundant data across reads or alignments, decreasing file size and allowing size to grow sub-linearly (rather than linearly) with the number of reads. CRAM (11), NGC (25), Goby (4) and REFEREE (7) use reference-based compression, which was proposed earlier (6, 16), to replace a read sequence with a concise description of how it differs from a substring of the reference. Quip (12) uses arithmetic coding together with a sequence model trained on-the-fly to compress losslessly and without a reference.

Goby uses a range of strategies, including column-wise compression and detailed modeling of relationships between columns. REFEREE uses separable streams and clustering of quality strings. In these formats, the alignments and the fields are largely preserved, but are compressed along with neighbors row-wise (together with the other fields of the same alignment), or column-wise (with other instances of the field across alignments).

These studies also explore lossy compression schemes, in which less important data, such as read names and quality strings, is selectively discarded. Many tools optionally discard read names and quality values, and REFEREE clusters quality strings and replaces each with a single representative from its cluster.

Boiler takes a radically lossy approach to compressing RNA-seq alignments, yielding very small compressed outputs. Inspired by the notion of transform coding, Boiler converts alignment data from the “alignment domain,” where location, shape and pairing information are stored for every alignment, to the “coverage domain,” where the coverage vector is stored and alignment information is inferred where needed. Boiler keeps only a set of coverage vectors and a few empirical distributions that partially preserve fields such as `POS` (offset into chromosome) `TLEN` (genomic outer distance), `XS:i` (strand) and `NH:i` (number of hits). Consequently, Boiler is lossy in an unusual sense: compressing and decompressing might cause alignments to shift along the genome, change shape, or become matched with the wrong mate. Table 1 presents a comparison of how CRAM, Goby, and Boiler preserve read information.

Boiler should not be considered a general-purpose alignment compression tool. Because it discards quality values and non-reference alleles, its output is not appropriate for downstream tools requiring such data, such as variant callers and tools for allele-specific expression. However, Boiler preserves the data most relevant to popular downstream RNA-seq tools for quantification, assembly and differential expression. We show that popular isoform-level tools — Cufflinks (28) and StringTie (24) - yield near-identical results when the input is Boiler-compressed.

Boiler yields extremely small file sizes, more than 3-fold smaller than files produced by CRAM and Goby for paired-end samples with at least 10M paired-end reads. Unlike other compression tools, Boiler’s compression ratio improvessubstantially as input file size grows, growing from about 10fold for lower-coverage unpaired samples to over 50-fold for higher-coverage samples. Speed and memory footprint are comparable to other compression tools despite the fact that, as we show, recovering alignments from a coverage vector is computationally hard. Also, because nucleotide data is removed, Boiler-compressed data is effectively de-identified, making it easier to pass between parties securely.

Boiler also provides a range of speedy queries. Many compression tools provide a way for the user to extract alignments spanning a particular genomic interval from the compressed file. REFEREE goes a step further by enabling faster queries when the user is concerned with only a subset of the fields. Boiler goes further still by providing fast queries that are directly relevant to downstream uses of RNA-seq data. Boiler allows the user and downstream tools to (a) iterate over “bundles” of alignments according to inferred gene boundaries, (b) extract the coverage vector across a genomic interval, and (c) extract alignments overlapping a genomic interval.

## MATERIALS AND METHODS

### Compression

Boiler implements a lossy compression scheme that preserves only the data needed by downstream isoform assembly tools such as Cufflinks and StringTie. For this reason, read names and quality strings are discarded, along with other data that has little or no bearing on downstream RNA-seq analysis.

Given a set of alignments to a reference genome, Boiler first partitions the alignments into “bundles” of overlapping reads. Bundles are computed in the same manner as Cufflinks’ initial bundling step: As sorted reads are processed, if the current read starts within 50 bases of the end of the current bundle, the read is added to the bundle. Otherwise, the current bundle is compressed and a new bundle is initialized beginning with the current read.

Boiler converts each bundle into a set of coverage vectors and tallies of observed read lengths. For most Illumina sequencing datasets, reads are uniform-length (or nearly so, e.g. after trimming), yielding a concise tally. If any alignments in the bundle are paired-end, Boiler also stores a tally of observed genomic outer distances as reported by the aligner in the `TLEN` SAM field. Note that `TLEN` includes the lengths of all the introns spanned by the alignment, so we refer to this as “genomic outer distance,” rather than “fragment length.” The coverage vector is compressed using run-length encoding, which is particularly effective in low-coverage regions.

**Table 1.** Comparison of the SAM fields stored by different compression tools. CRAM and Goby can preserve some fields through configurable options, summarized in the “Config” columns.

| SAM Feature | CRAM |  | Goby |  | Boiler |
| --- | --- | --- | --- | --- | --- |
|  | Default | Config. | Default | Config. |  |
| Read Name | Yes<sup>1</sup> | No | No | Yes | No |
| Flags | Yes | No | Yes | No | No |
| Mapping Quality | Yes | No | Yes | No | No |
| Read Sequence | Yes | No | Yes | No | No |
| Quality Scores | No | Yes | Yes<sup>2</sup> | Yes | No |
| Tags | No | Yes | MD | Yes<sup>3</sup> | XS, NH |

<sup>2</sup>For mismatches only
<sup>3</sup>Goby preserves either all tags or just the MD tag.

^1^CRAMtools documentation claims that by default read names should not be preserved, however we were not able to replicate this functionality.

^2^For mismatches only

^3^Goby preserves either all tags or just the MD tag.

Each bundle is compressed as follows:

1. Boiler scans the bundle’s alignments to find splice sites spanned by at least one alignment. Boiler divides the portion of the genome spanned by the bundle into “partitions” formed by cutting at every splice site (Figure 1a).
2. Boiler assigns each alignment to a *bucket* according to: (a) the subset of partitions spanned by the alignment, (b) the value in the alignment’s `NH:i` field, indicating the number of distinct locations where the read aligned to the reference, and (c) the value in the `XS:A` field, indicating whether spanned splice motifs are consistent with the sense (+) or anti-sense (-) strand of the gene. Alignments not spanning a junction usually lack the `XS:A` field; Boiler treats these as though the `XS:A` field contains a “dummy” value indicating the strand is unknown.
3. For each bucket, Boiler computes the coverage vector from the alignments assigned to it. Boiler writes the run-length encoded coverage vector (Figure 1b) followed by the read and genomic outer distance distributions (Figure 1c) for the junction.

Each bundle, which consists of many buckets, is compressed independently using the DEFLATE algorithm as implemented in the `zlib` package from the Python Standard Library. Each bundle is compressed separately to make targeted queries efficient, as discussed in the “Queries” section.

Some RNA-seq alignment tools (including HISAT (15)) output SAM records for reads or ends that fail to align, whereas others (including TopHat 2 (14)) do not. Boiler deals only with aligned reads. SAM records for unaligned reads are ignored, and those reads are not represented in a compressed Boiler file. Additionally, if one end of a paired-end read is “orphaned” — i.e. its opposite end fails to align — Boiler will convert the orphan to an unpaired read. The paired nature of orphaned reads is lost during Boiler compression.

### Unbundled alignments

Prior to compression, Boiler must identify and handle paired-end alignments that span bundles in unexpected ways. We call these *unbundled* alignments. Unbundled alignments fall into four categories: (a) one end falls within an intron spanned by the other end, (b) the two ends align to different chromosomes, (c) the two ends align to the same chromosome but very far from each other, (d) one end is assigned to the sense strand, while its mate is assigned to the anti-sense strand. Both TopHat and HISAT report such alignments, though they constitute only a small fraction of the alignments in a typical dataset.

These alignments are hard to fit into the bundling scheme described previously. Reads in category (a) are biologically implausible. Boiler treats them as unpaired reads by default, however the user may choose to preserve these pairings.

**Figure 1.**
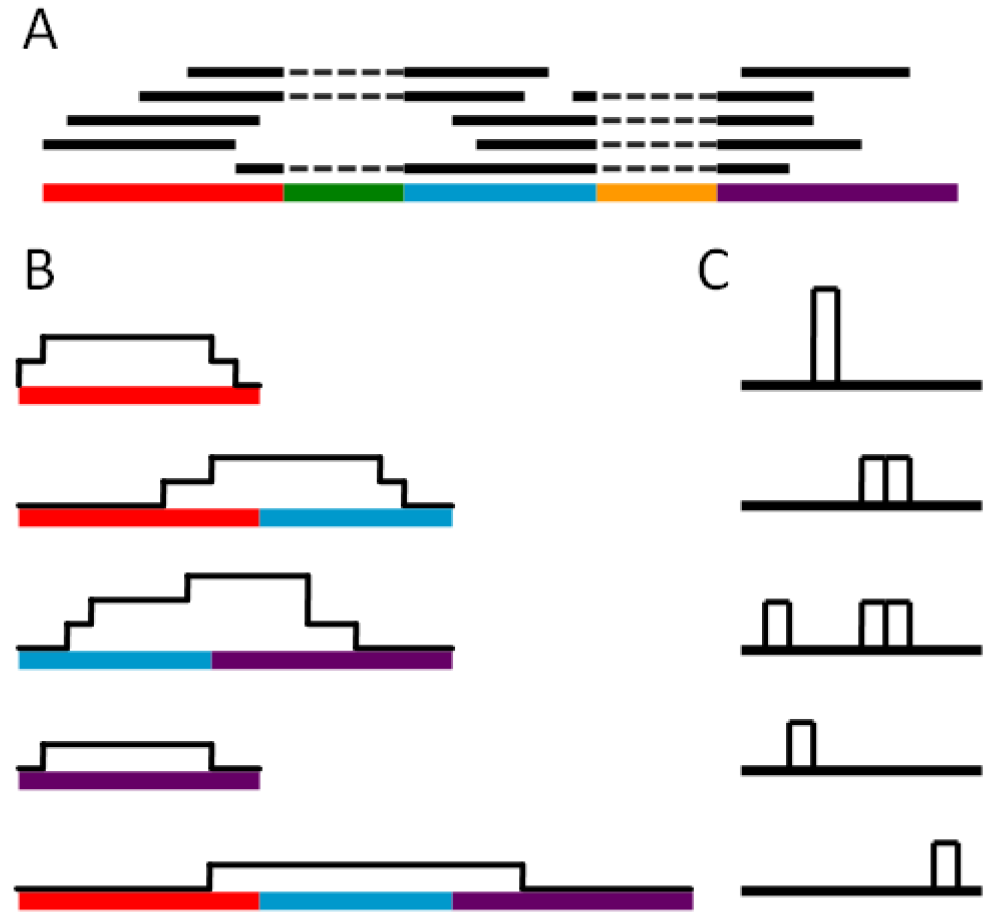
Illustration of how Boiler compresses alignments in a bundle, for a dataset with unpaired reads. (a) The genome is divided into “partitions” (colored segments) based on the processed splice sites. A bucket is defined by the subset of partitions spanned (as well as the values of the NH:i and XS:A fields, though these are omitted from the figure for simplicity). Each bucket stores (b) the coverage vector and (c) the length tally of the reads assigned to the bucket.

Categories (b) and (c) could be scientifically relevant and should be preserved. For instance, alignments in category (b) may be evidence of gene fusions. Boiler stores all alignments in categories (b) and (c) in a special “unbundled alignments” section of the compressed file. Unbundled alignments are stored in *bundle-spanning buckets*. A bundle-spanning bucket is identical to a normal bucket, but includes the indices of the two bundles it spans in addition to the list of partitions spanned from each bundle. The bundle-spanning buckets are stored as a contiguous list, compressed in small chunks using zlib, and indexed to reduce work for targeted queries.

Treatment of pairs in category (d) is configurable by the user. By default, they remain paired and one end, selected at random, is modified to match the other’s strand. Optionally (using ——`split`—`discordant`), such pairs can be treated as unpaired reads, which is consistent with how they are treated by Cufflinks and StringTie.

### Multi-mapping reads

RNA-seq reads may align equally well to many genomic locations. Such reads are called “multi mappers.” Because downstream tools might treat multi mappers differently from uniquely mapped reads, at least some multi-mapping information must be preserved by Boiler.

For a multi-mapping read, Boiler preserves each alignment along with its NH:i extra field. However, Boiler also discards read names. This can have an adverse impact on downstream tools that rely on read names to establish the one-to-many relationship between reads and their alignments. Our fidelity experiments show this does not substantially impact the accuracy of downstream tools for isoform assembly and quantification. However, it does have an adverse effect on quantification accuracy when Cufflinks is asked to quantify from a given gene annotation (`-G` mode), as discussed in Supplementary Note 11. StringTie’s accuracy when quantifying from a given gene annotation (`-G -e` mode) is not substantially affected.

### Decompression

To decompress, Boiler first expands each bundle with the INFLATE algorithm as implemented in the Python zlib module, then expands each bucket.

When decompressing a bucket, Boiler seeks to recreate the set of alignment intervals that yielded the bucket’s coverage vector and read and genomic outer distance tallies. This is a two-step process; first reads must be recovered from the coverage vector and read length tally (“read recovery”), then the recovered reads must be paired according to the paired length tally (“pairing”).

The *read recovery* problem may not have a unique solution; e.g., consider a compressed dataset with read lengths *l*_1_ and *l*_2_ (*l*_1_ ≠ *l*_2_) and a coverage vector containing 1 at all positions in the range [0, *l*_2_ + *l*_2_). This case has two valid solutions:

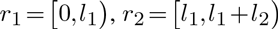

and

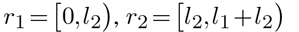

Thus, we cannot guarantee perfect recovery of the compressed reads.

We define the read recovery problem as follows. Given a coverage vector and tally of read lengths, we seek a list of decompressed reads (genomic intervals) such that

1. The decompressed read lengths are a subset of those given in the tally,
2. at no position does the coverage vector produced by the decompressed reads exceed the value in the original coverage vector, and
3. the sum of the lengths of all decompressed reads is maximized.

This formulation is general enough to tolerate an input where the read length tally and coverage vector are not compatible, i.e., where no solution fits both precisely. In this case, the algorithm might decompress only some of the reads in the input tally.

We observe that the read recovery problem is NP-hard in general (proved by reduction from the Multiple Subset Sum Problem in Supplementary Note 1), but that some special cases are easily solved. When all reads are the same length, for example, the solution is unique and can be found efficiently. We also observe that second-generation sequencing produces datasets with uniform or near-uniform (e.g. after trimming) read-length tallies. These facts lead us to propose the greedy algorithm described below. The algorithm is not optimal in general, but it is well suited to cases where the input read lengths are uniform or almost uniform.

*Greedy algorithms for extracting and pairing reads*. The algorithm works from one end of the coverage vector to the other, extracting reads that are “consistent” with coverage. A read is consistent with coverage if removing the read and decrementing the corresponding coverage-vector elements does not cause any vector elements to fall below zero. When a consistent read is selected for extraction, the corresponding coverage-vector elements are decremented and the process repeats. The process stops when the far end of the coverage vector is reached.

When reads have uniform length, the algorithm yields the correct solution. When reads have various lengths, the problem is harder and the algorithm may fail to yield the optimal solution. In the case where reads are various lengths, Boiler’s algorithm uses heuristics to arrive at a solution where (a) the coverage vector induced by the extracted reads matches the true vector as closely as possible, and (b) the distribution of extracted read lengths matches the true distribution of read lengths as closely as possible. Boiler favors (a) over (b); i.e. it will artificially lengthen or shorten the extracted reads to fix small coverage discrepancies. The heuristics are described in Supplementary Note 2.

If the data contains paired-end reads, we must also solve the *read pairing* problem. We would like Boiler to restore the paired-end relationships in a way that matches the original genomic outer distances as closely as possible. We start with (a) a collection of reads (ends), all of which are initially unpaired, and (b) the “true” genomic outer distance tally, storing the frequencies of each distance, which was compiled and stored during compression.

We use the following greedy algorithm to pair up reads in a way that closely matches the true genomic outer distance tally. Each read is examined, working inward from the extremes, alternating between the left and right extremes. For each read, we seek the most distant read such that the resulting pairing is compatible with distances remaining in the tally. When two reads are paired in this way, they are removed from future consideration, and the corresponding element of the tally is decremented. Reads that are not paired in this way are matched up randomly in a second pass.

### Queries

Boiler allows the user to query a compressed RNA-seq dataset to (a) iterate over genomic intervals delimiting regions of non-zero coverage, roughly corresponding to genes, (b) extract the genomic coverage vector across a specified genomic interval, and (c) extract alignments overlapping a specified genomic interval. Because each bundle of alignments is compressed separately, Boiler can answer such queries without decompressing the entire file.

Bundle boundaries are stored in the index at the beginning of the compressed file, so skipping to a particular bundle can be accomplished with a single uncompressed index lookup. To compute the coverage query (query *b*, above), Boiler combines the relevant portions of the coverage vectors for all the buckets overlapping the specified region. This requires that Boiler decompress the DEFLATED and run-length encoded coverage vectors, but does not require the more work-intensive read and pair recovery algorithms. Alignment-level queries (queries *a* and *c* above) are more expensive, requiring Boiler to run the greedy read recovery algorithm on each of the buckets overlapping the specified region.

Downstream tools like Cufflinks can be modified to query Boiler-compressed files directly, removing the need for an intermediate SAM/BAM file. When a downstream tool requires access only to information about gene boundaries (query *a*, above) or about targeted regions of the coverage vector (query *b*), Boiler’s queries can be much faster than directly querying a sorted and indexed BAM file.

### Implementation

Boiler is implemented in Python and is compatible with Python interpreters version 3 and above. All of the Python modules used by Boiler are in the Python Standard Library, making Boiler quite portable across Python installations and interpreters. For example, we use the fast PyPy interpreter for our experiments.

## RESULTS

We used Flux Simulator v1.2.1 (19) to simulate 10 RNA-seq samples from the BDGP5 build of the *D. melanogaster* genome and the Ensembl release 70 (29) gene annotation. We simulated both paired-end and unpaired RNA-seq samples for a series of library sizes: 0.5, 1, 2.5, 5, 10, and 20 million reads. We also simulated two samples from the hg19 build of the human genome and Gencode v12 gene annotation (10) containing 20 and 40 million paired-end reads. We also used two real human RNA-seq samples. We used sample HG00100 from the GEUVADIS (17) study, consisting of about 20 million paired-end reads. We also chose one of the seven technical replicates of the 3:1 ratio of Universal Human Reference RNA to Human Brain Reference RNA from the SEQC study (26). The study accession is SRP025982 and the individual replicates have run accessions SRR1216073 - SRR1216079. Each replicate consists of approximately 11- million paired-end reads. We tested sample SRR1216073, labeled “SRP025982” in the results below. All samples were aligned to the reference genome using either TopHat 2 v2.1.0 (14) with default parameters, or HISAT 0.1.6-beta (15) with default parameters. *D. melanogaster* samples were aligned to the BDGP5 reference and human samples were aligned to the hg19 reference.

We compared Boiler’s speed, compression ratio, and peak memory usage to Goby and CRAMTools. Boiler and Goby remove read names by default, but CRAM does not. (CRAMtools has an option to preserve read names, but we cannot find a working mechanism in version 3 to remove them.) For a fair comparison, we stripped the read names before compressing. For Goby, we enabled the full”ACT H+T+D” compression scheme, as described in the Goby study (4).

Further details on software versions and command-line arguments used are included in Supplementary Note 3.

### Efficiency and compression ratio

We compressed each TopHat 2 alignment file with Boiler v1.0.1, CRAMTools v3.0, and Goby v2.3.5. Boiler was run with PyPy v2.4 and CRAMTools and Goby were run with Java v1.7. All tools were run on the Homewood High
Performance Compute Cluster at Johns Hopkins University. Each cluster computer has 2 Intel Xeon X5660 2.80GHz processors and 48 GB of RAM. We measured running time by adding the user and sys times reported by the Linux time command. Each tool runs predominantly on a single thread and processor. We measured peak memory usage in Python by spawning a new child process for the command and polling maximum resident set size (RSS) using the Python resource package’s getrusage function. Peak memory usage for Boiler and Goby was consistent across runs, but CRAM memory usage varied widely between runs. We report the median peak memory of 10 runs for greater consistency.

Boiler takes roughly 1.5-2 times longer than CRAMTools and Goby to compress the D. melanogaster samples and about 2-12 times longer for the human samples (Table 2, Figure 2). It requires less memory than Goby for all but the deeper human samples, and less than CRAM for small datasets. For larger datasets, CRAM memory’s memory footprint seems to be capped at around 2 GB (Table 3). Supplementary Note 5 compares the decompression time for all three tools.

Importantly, Boiler achieves a compression ratio comparable to BigWig (Table 4) for all but the largest paired-end datasets, and usually produces far smaller compressed files than CRAMTools or Goby. We measured both compressed file size (Table 4) and the “compression ratio” of original to compressed file size (Figure 3) for alignments generated by TopHat 2. The “original” file is a sorted BAM file with read names removed. For low-coverage unpaired datasets, CRAM and Goby’s compression ratios are superior to Boiler’s. However, we observe that while CRAMTools and Goby’s compression ratios remain flat as the *D. melanogaster* library size increases, Boiler’s ratios improve substantially (Figure 3), achieving its best compression ratios for the 20M-read samples: 56-fold for unpaired and 39fold for paired-end samples. Boiler’s compression ratio is consistently better than the other tools for paired-end samples and improves as library size increases. For high coverage *D. melanogaster* and all human datasets, Boiler achieves compression ratios 3-5-fold higher than both CRAM and Goby.

To demonstrate Boiler’s performance on an established benchmark, we also ran Boiler on the dataset with Sample ID EJOYQAZ from the Goby study (Table 4). This consists of roughly 7.5 million paired-end reads from *H. sapiens*. The BAM file as well as the compressed Goby output is available at data.campagnelab.org/home/compression-of-structured-high-throughput-sequencing-data. Boiler produces a compressed file of 26.9 MB (2.8% of the original BAM) compared to Goby’s output of 123.2 MB (13.0% of the original BAM), representing a 77% space reduction for Boiler compared to Goby.

Boiler achieves a comparable compression ratio for alignments generated by HISAT (Supplementary Note 4). We also show in Supplementary Note 4 that removing orphaned reads or quality scores from the SAM file does not significantly impact Goby’s compression ratio.

**Figure 2.**
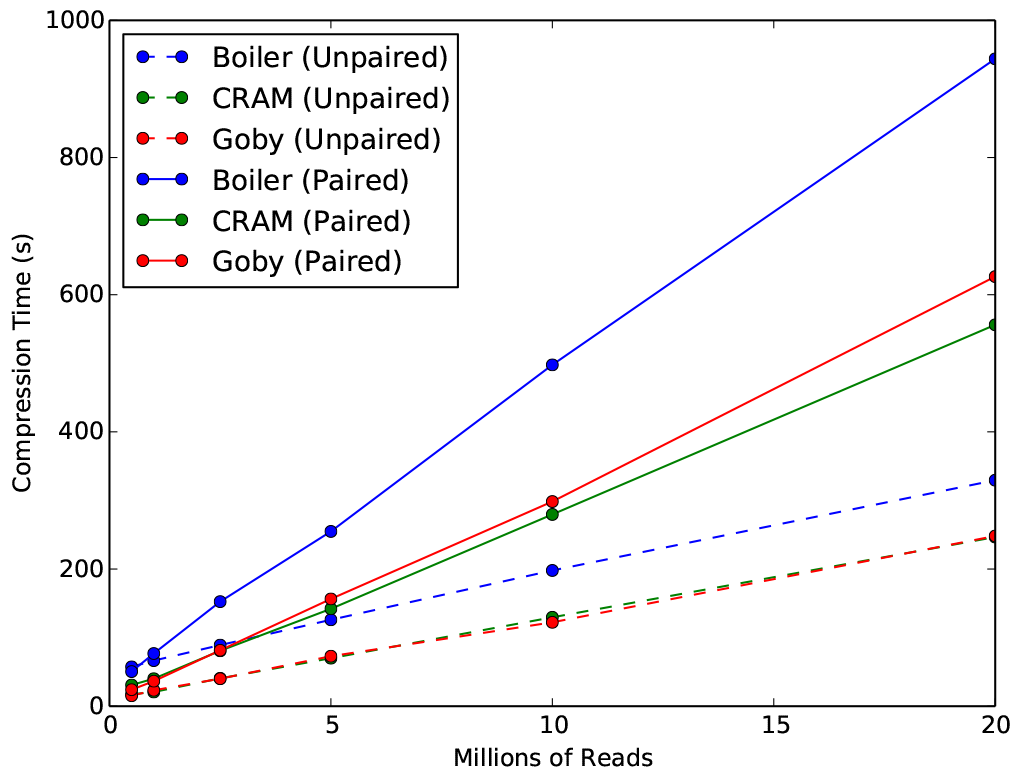
Time required for Boiler, CRAM and Goby to compress simulated *D. melanogaster* paired-end datasets.

### Fidelity

Boiler discards read nucleotide and quality-value data. So while Boiler is not appropriate for pipelines where downstream tools measure non-reference alleles — e.g. for variant calling, allele-specific expression, or RNA editing — we show that Boiler is appropriate for the common case where downstream tools are concerned with assembling and quantifying isoforms, e.g. Cufflinks and StringTie.

Boiler tends to “shuffle” alignment data in certain ways during compression. Some of the shuffling is harmless, having no adverse effect on downstream results from Cufflinks and StringTie. But some shuffling could be harmful, negatively impacting the fidelity of downstream results. Using both simulated and real data, we (a) establish the nature of the shuffling introduced by Boiler, (b) show there is only slight harmful shuffling in practice, and (c) show that the overall amount of shuffling is smaller - often much smaller - than the shuffling that results from substituting one technical replicate for another.

**Table 2.** Compression times in seconds.

| Dataset | Boiler | CRAM | Goby |
| --- | --- | --- | --- |
| Drosophila, Simulated Unpaired |  |  |  |
| 0.5M | 57.4 | 17.4 | 15.3 |
| 1M | 66.5 | 20.7 | 23.2 |
| 2.5M | 89.1 | 40.6 | 40.1 |
| 5M | 126.0 | 70.0 | 72.9 |
| 10M | 197.9 | 129.6 | 122.3 |
| 20M | 329.5 | 246.6 | 248.2 |
| Drosophila, Simulated Paired |  |  |  |
| 0.5M | 50.5 | 30.9 | 24.1 |
| 1M | 76.8 | 40.1 | 36.5 |
| 2.5M | 152.5 | 80.8 | 81.5 |
| 5M | 254.8 | 142.0 | 156.4 |
| 10M | 497.7 | 279.6 | 298.4 |
| 20M | 943.9 | 556.1 | 626.4 |
| Human |  |  |  |
| SRP025982 (11M) | 6122.6 | 507.6 | 674.5 |
| HG00100 (20M) | 4118.8 | 691.6 | 1045.6 |
| Simulated 20M | 2337.3 | 906.8 | 1301.3 |
| Simulated 40M | 7906.3 | 1476.8 | 2138.7 |

### Alignment-level fidelity

Boiler compression can change where alignments lie on the genome and how they are paired. Here we ask how well alignment locations are preserved after Boiler compression of the TopHat 2-aligned samples. We measure alignment-level precision and recall in two ways. First we ignore read pairings. For each aligned unpaired read (or end of a paired-end read) in the original file, we seek a corresponding alignment in the compressed file where the genomic position of the alignment and of all overlapped splice junctions are identical. This counts as a true positive, and the alignments involved are “matched.” An alignment can be matched with at most one other alignment. An alignment in the original file that fails to match an alignment in the compressed file counts as a false negative and the converse is a false positive Given these definitions, precision and recall are shown in the left-hand columns of Table 5 (labeled “Ignoring pairings”). These range from 96.2% to 99.4% across the samples tested.

**Table 3.** Peak memory usage (GB) reported by Python. Numbers reported are the median across 10 runs.

| Dataset | Boiler | CRAM | Goby |
| --- | --- | --- | --- |
| Drosophila, Simulated Unpaired |  |  |  |
| 0.5M | 0.38 | 0.74 | 0.73 |
| 1M | 0.29 | 1.37 | 1.39 |
| 2.5M | 0.24 | 1.12 | 1.72 |
| 5M | 0.24 | 1.13 | 1.73 |
| 10M | 0.24 | 0.96 | 2.11 |
| 20M | 0.50 | 1.53 | 3.77 |
| Drosophila, Simulated Paired |  |  |  |
| 0.5M | 0.32 | 1.15 | 0.90 |
| 1M | 0.31 | 1.91 | 1.05 |
| 2.5M | 0.30 | 1.96 | 1.33 |
| 5M | 0.32 | 1.18 | 1.61 |
| 10M | 0.79 | 1.11 | 5.67 |
| 20M | 1.57 | 0.99 | 7.21 |
| Human |  |  |  |
| SRP025982 (11M) | 7.10 | 2.08 | 5.3 |
| HG00100 (20M) | 3.23 | 2.08 | 4.12 |
| Simulated 20M | 6.11 | 2.08 | 4.87 |
| Simulated 40M | 9.14 | 2.08 | 6.88 |

**Table 4.** Size of compressed files (MB) compared to the original sorted BAM with read names removed.

| Dataset | BAM | BigWig | Boiler | CRAM | Goby |
| --- | --- | --- | --- | --- | --- |
| Drosophila, Simulated Unpaired |  |  |  |  |  |
| 0.5M | 26.0 | 3.7 | 2.7 | 0.9 | 1.2 |
| 1M | 49.4 | 6.4 | 3.6 | 1.6 | 2.4 |
| 2.5M | 120.3 | 12.8 | 5.6 | 3.8 | 5.6 |
| 5M | 222.2 | 19.0 | 7.6 | 6.7 | 10.1 |
| 10M | 436.1 | 28.1 | 11.0 | 12.6 | 19.0 |
| 20M | 883.0 | 37.0 | 15.0 | 23.2 | 35.7 |
| Drosophila, Simulated Paired |  |  |  |  |  |
| 0.5M | 56.1 | 6.3 | 4.3 | 7.7 | 4.9 |
| 1M | 104.6 | 10.4 | 6.7 | 14.2 | 9.4 |
| 2.5M | 255.6 | 19.8 | 12.9 | 33.9 | 23.8 |
| 5M | 488.1 | 27.8 | 20.0 | 61.9 | 45.9 |
| 10M | 955.0 | 37.2 | 31.6 | 116.9 | 95.8 |
| 20M | 1902.6 | 47.8 | 49.0 | 226.2 | 193.3 |
| Human |  |  |  |  |  |
| EJOYQAZ (7.5M) | 747.9<sup>1</sup> | – | 26.9 | – | 123.2<sup>1</sup> |
| SRP025982 (11M) | 810.1 | 54.6 | 38.0 | 160.5 | 159.8 |
| HG00100 (20M) | 2017.9 | 98.7 | 78.0 | 288.8 | 352.2 |
| Simulated 20M | 2117.3 | 54.3 | 71.6 | 273.5 | 307.8 |
| Simulated 40M | 3858.4 | 71.7 | 118.0 | 491.4 | 587.6 |

<sup>1</sup> Datasets downloaded from Goby supplementary data.

^1^Datasets downloaded from Goby supplementary data.

**Figure 3.**
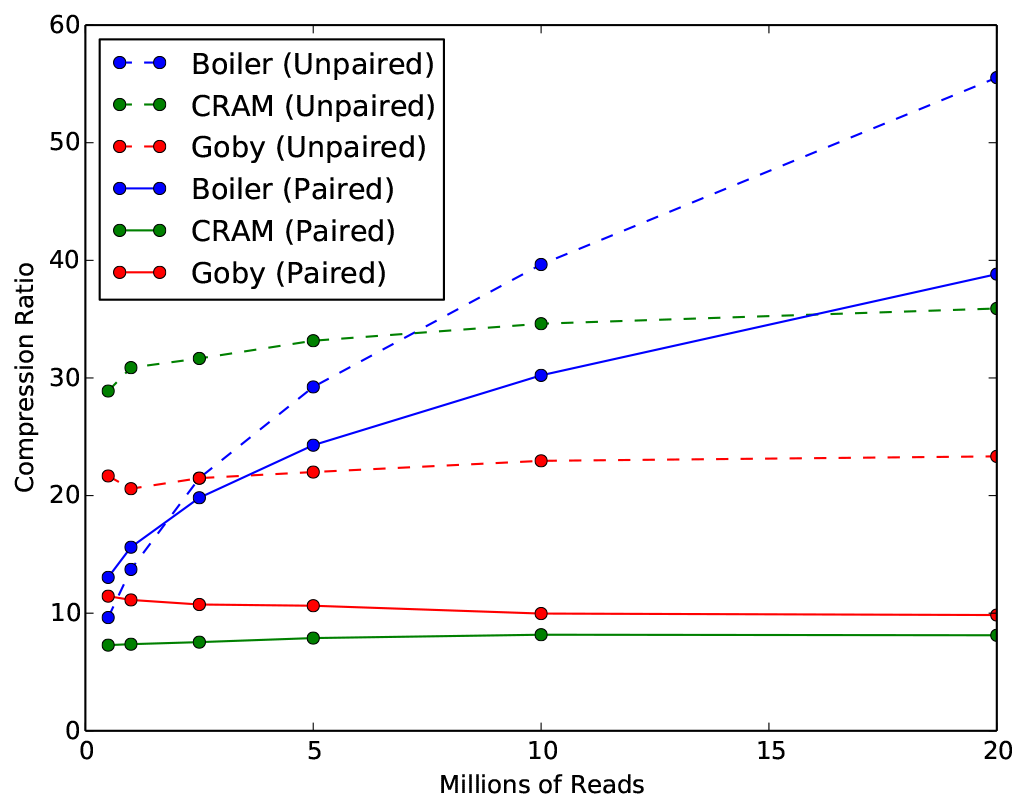
Compression ratios for simulated *D. melanogaster* datasets when compressed by Boiler, CRAM and Goby. Ratios are with respect to the original sorted BAM with read names removed.

We also measure precision and recall in a way that takes pairing into account: for each aligned pair, we seek a corresponding pair in the compressed file where both ends match their counterparts in terms of their genomic position and the positions of splice junctions. These results are shown in the right-hand columns of Table 5 (labeled “Including pairings”). Here precision and recall are lower, with most samples ranging from 17.6% to 54.5%. Note that the SEQC sample, SRP025982, exhibited higher unpaired precision and recall than the others (99.4%), and much higher than the others when considering pairing (89.4%). This is likely due to the smaller number of reads in the sample relative to the other human samples, and to the much larger number of Boiler buckets induced by that sample. The larger number of buckets is likely owing to the presence of Universal Human Reference RNA, in which many genes are expressed.

We also measure genomic outer distance distribution (excluding unbundled alignments) before and after Boiler compression. We find they match closely (Figure 4, left) though not perfectly (Figure 4, right). The results are emblematic of Boiler’s strategy: aggregate distributions are preserved, but links between particular alignments and particular points in the distribution are lost. As a result, somedata is “shuffled;” ends themselves are largely unchanged, but pairings between ends are shuffled in a way that preserves the aggregate genomic outer distance distribution.

We repeated these experiments for alignments output by HISAT, as discussed in Supplementary Note 7. The results are similar to those produced by TopHat 2, with precision and recall ranging from 96.7% to 99.3% when ignoring pairings, and from 14% to 32.1% when considering pairings.

### Isoform fidelity

Having established Boiler’s lossy-ness and shuffling behavior, we now assess the degree to which loss and shuffling have an adverse effect on downstream results obtained by Cufflinks v2.2.1 and StringTie v1.2.2. StringTie was run with default parameters. Cufflinks was run with the `——no—effective—length—correction` parameter to avoid variability due to an issue (recently resolved) in how Cufflinks performs effective transcript length correction (5).

Let *T* be the true simulated transcriptome, including abundances for each transcript, which we extract from the Flux-generated .`pro` and .`gtf` files. Let 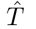 be the transcriptome assembled and quantified from the original alignments, which we extract from the Cufflinks/StringTie output. 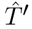 be the same but for the Boiler-compressed alignments. Here we ask whether 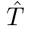 and 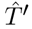 are approximately equidistant from *T*, indicating Boiler’s loss and shuffling are not having an adverse effect.

We define a function for measuring the distance between two transcripts *t*_1_ and *t*_2_ assembled with respect to a reference genome. The function outputs a value between 0 and 1, with 0 indicating the transcripts do not match and 1 indicating a perfect match.

A transcript *t* can be represented as a set of exons {*e*_1_,…*e_n_*}, each defined by its start and stop positions. We first define a scoring function for two exons *e*_1_ = (*x*_1_,*y*_1_) and *e*^2^ = (*x*^2^,*y*^2^):

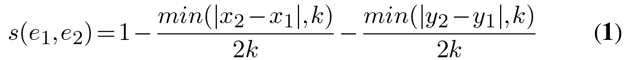

for some threshold *k*. We further define function 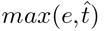 to be the set Ê of exons ê from 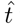 with maximal score *s(e,ê)*.

**Table 5.** Precision and recall of SAM reads.

| Dataset | Ignoring Pairings |  | Including Pairings |  |
| --- | --- | --- | --- | --- |
|  | Precision | Recall | Precision | Recall |
| Drosophila, Simulated Unpaired |  |  |  |  |
| 0.5M | 0.991 | 0.993 | – | – |
| 1M | 0.986 | 0.989 | – | – |
| 2.5M | 0.976 | 0.981 | – | – |
| 5M | 0.969 | 0.974 | – | – |
| 10M | 0.962 | 0.968 | – | – |
| 20M | 0.964 | 0.969 | – | – |
| Drosophila, Simulated Paired |  |  |  |  |
| 0.5M | 0.990 | 0.992 | 0.544 | 0.545 |
| 1M | 0.984 | 0.987 | 0.458 | 0.460 |
| 2.5M | 0.972 | 0.978 | 0.338 | 0.340 |
| 5M | 0.967 | 0.972 | 0.263 | 0.264 |
| 10M | 0.964 | 0.969 | 0.219 | 0.220 |
| 20M | 0.964 | 0.968 | 0.176 | 0.177 |
| Human |  |  |  |  |
| SRP025982 (11M) | 0.994 | 0.994 | 0.894 | 0.894 |
| HG00100 (20M) | 0.983 | 0.983 | 0.261 | 0.261 |
| Simulated 20M | 0.973 | 0.976 | 0.327 | 0.328 |
| Simulated 40M | 0.972 | 0.975 | 0.319 | 0.320 |

**Algorithm 1.**
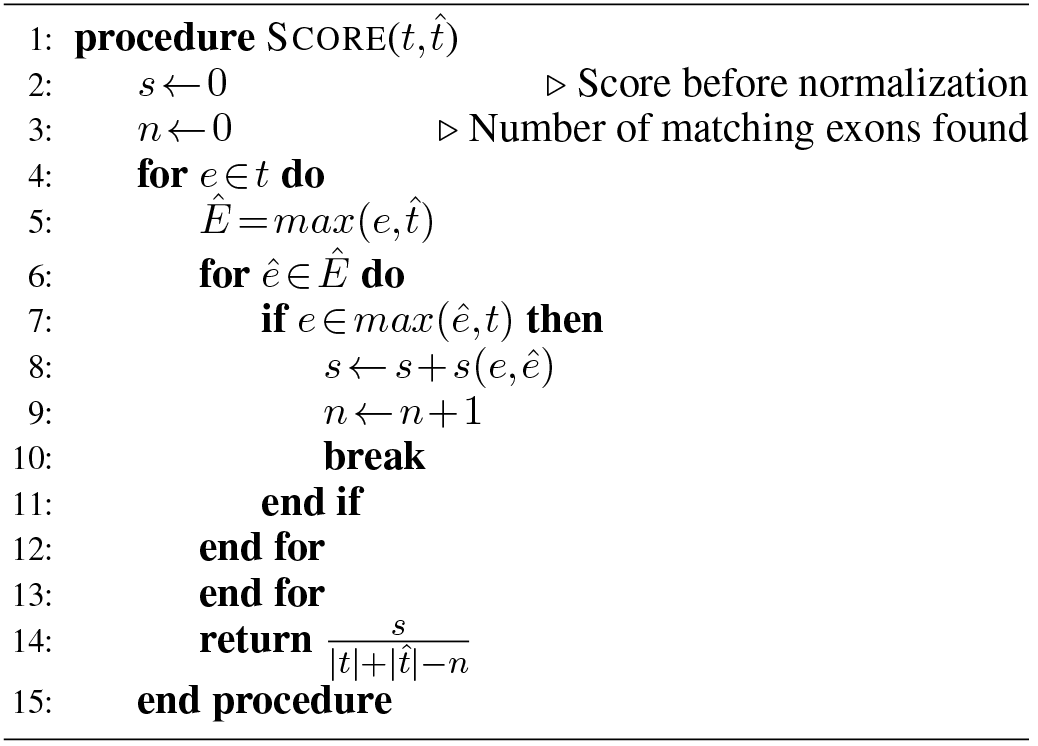
Transcript scoring algorithm

The weighted precision for pre-compression alignments is:

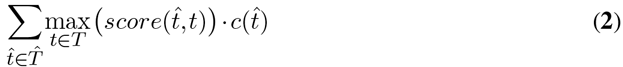

Where 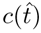 is the predicted coverage level of *t* as reported by Cufflinks/StringTie. This measure is weighted both by the coverage of the assembled transcript and by the similarity of the matched-up transcripts. Precision for the post-compression alignments is calculated similarly, using 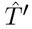 instead of 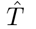.

Similarly, weighted recall is

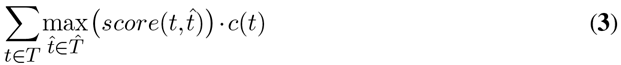

Where *c(t)* is the true coverage level for *t* as reported by Flux.

We used a threshold of *k* = 10 for the exon scoring function (1). The selection of *k* = 10 is discussed in Supplementary Note 6. We calculated the weighted precision (Tables 6) and recall (Table 7) for all simulated samples. While there are small differences in all experiments (as expected due to Boiler’s shuffling behavior) overall Boiler compression does not have a substantial adverse impact on weighted precision and recall.

### Other Isoform Fidelity Scores

Supplementary Note 9 describes a complementary experiment using weighted k-mer recall (WKR), which does not depend on a distance function. Supplementary Note 10 describes a third accuracy measure called the Tripartite Score, which uses the same distance, but can compare two assembled transcriptomes to a third “reference” transcriptome. The results from those methods substantially agree with the results above.

**Figure 4.**
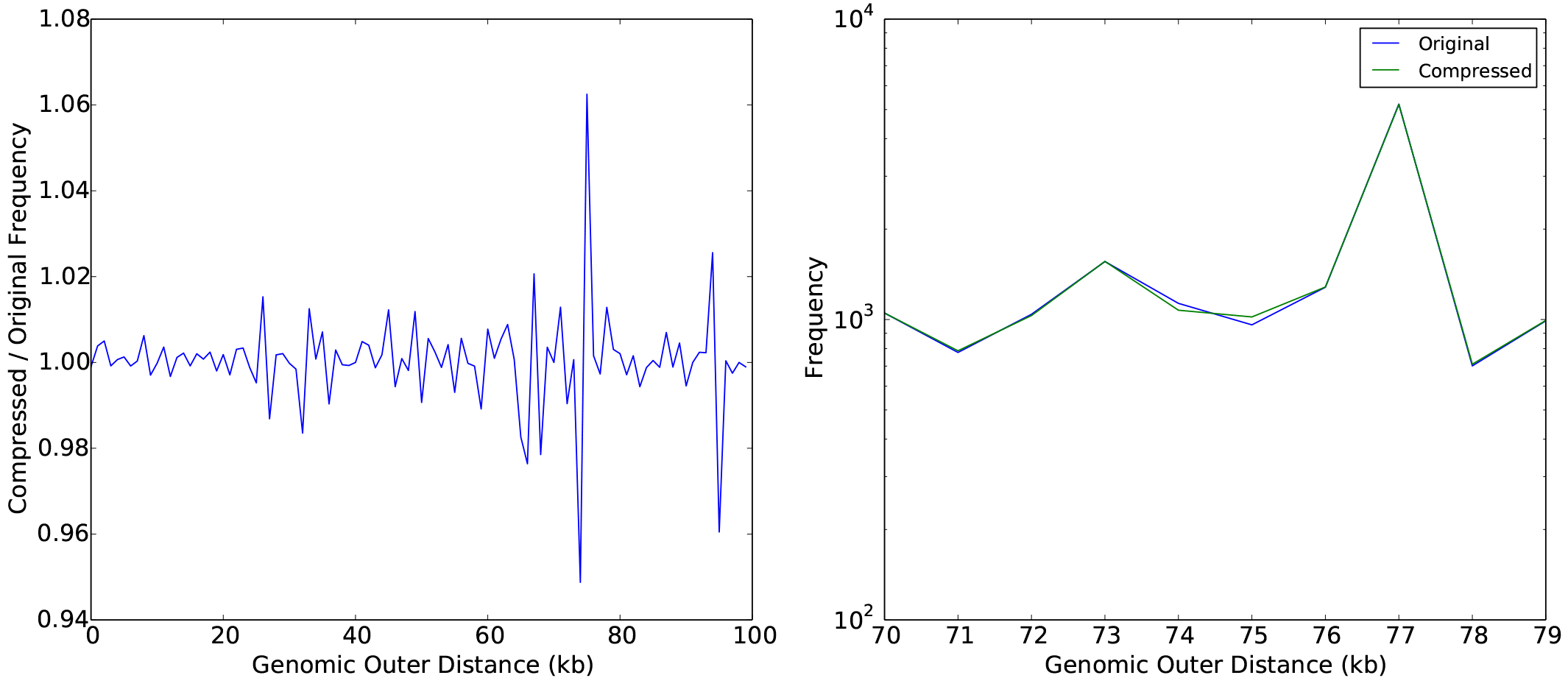
Comparison of genomic outer distances for the 10M paired-end *D. melanogaster* sample. Left: Frequency ratio of each genomics outer distance, compressed divided by original, up to a distance of 100,000 bases. Right: Original and compressed genomic outer distances between 70 and 79 kilobases in length.

### Shuffling relative to technical replicates

Next we investigate the amount of “shuffling” introduced by Boiler — causing some reads to shift along the genome and scrambling some paired-end relationships — and whether the effect is large or small compared to the shuffling that occurs when switching from one technical replicate to another.

**Table 6.** Reference-based Precision.

| Dataset | TopHat + Cufflinks |  | TopHat + StringTie |  | HISAT + StringTie |  |
| --- | --- | --- | --- | --- | --- | --- |
|  | Original | Compressed | Original | Compressed | Original | Compressed |
| Drosophila, Simulated Unpaired |  |  |  |  |  |  |
| 0.5M | 0.364 | 0.364 (+0.1%) | 0.466 | 0.466 (+0.0%) | 0.542 | 0.542 (+0.0%) |
| 1M | 0.432 | 0.434 (+0.3%) | 0.545 | 0.545 (+0.0%) | 0.611 | 0.611 (+0.0%) |
| 2.5M | 0.527 | 0.527 (+0.1%) | 0.627 | 0.627 (+0.0%) | 0.646 | 0.542 (-0.7%) |
| 5M | 0.564 | 0.564 (+0.1%) | 0.637 | 0.637 (-0.0%) | 0.644 | 0.641 (-0.5%) |
| 10M | 0.583 | 0.585 (+0.2%) | 0.645 | 0.645 (+0.0%) | 0.639 | 0.637 (-0.3%) |
| 20M | 0.602 | 0.603 (+0.2%) | 0.654 | 0.653 (-0.1%) | 0.642 | 0.638 (-0.6%) |
| Drosophila, Simulated Paired |  |  |  |  |  |  |
| 0.5M | 0.583 | 0.582 (-0.1%) | 0.546 | 0.546 (-0.0%) | 0.628 | 0.628 (-0.0%) |
| 1M | 0.611 | 0.609 (-0.3%) | 0.614 | 0.613 (-0.1%) | 0.660 | 0.660 (-0.0%) |
| 2.5M | 0.632 | 0.630 (-0.4%) | 0.644 | 0.643 (-0.2%) | 0.666 | 0.665 (-0.2%) |
| 5M | 0.635 | 0.633 (-0.3%) | 0.652 | 0.652 (-0.1%) | 0.662 | 0.661 (-0.2%) |
| 10M | 0.644 | 0.643 (-0.1%) | 0.663 | 0.662 (-0.1%) | 0.666 | 0.665 (-0.1%) |
| 20M | 0.639 | 0.634 (-0.8%) | 0.660 | 0.659 (-0.2%) | 0.660 | 0.657 (-0.4%) |
| Human, Simulated Paired |  |  |  |  |  |  |
| 20M | 0.552 | 0.554 (+0.4%) | 0.570 | 0.568 (-0.4%) | 0.613 | 0.613 (-0.1%) |
| 40M | 0.554 | 0.555 (+0.2%) | 0.576 | 0.572 (-0.6%) | 0.614 | 0.615 (+0.1%) |

We first construct five artificial technical replicates by generating five times the desired number of reads with Flux Simulator and randomly partitioning the resulting read file five ways. We then assemble and quantify each using Cufflinks and StringTie. Let 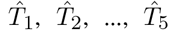 be the corresponding transcriptomes. We also pick a technical replicate (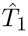, say) to compress with Boiler. Let 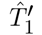 be the result of running Cufflinks/StringTie on the Boiler-compressed alignments. We calculate the weighted precision and recall of 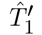 relative to 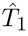. Finally, we calculate weighted precisions and recalls between all 10 ordered pairs of technical-replicate transcriptomes: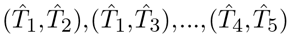. Results are presented in Tables 8 and 9. Precision and recall between technical replicates is shown as a range from the minimum to the maximum observed among the 10 ordered pairs. Precision and recall after Boiler-compression are consistently higher than precision and recall between technical replicates, indicating that shuffling due to Boiler compression is less consequential than shuffling due to technical variation.

For SRP025982, we followed the same process but using real lane-level technical replicates. The same sample was sequenced in 5 separate lanes of an Illumina instrument, but on the same flowcell. We compressed the first replicate (SRR1216073) with Boiler and used the four others (SRR1216076 - SRR1216079) to calculate the ten pairwise precision and recall measures.

It is notable how precision and recall change relative to per-sample coverage. Precision and recall between technical replicates increases as per-sample coverage increases, indicating that the shuffling effect decreases as more transcripts become deeply covered. On the other hand, precision and recall of 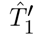 versus 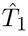 decreases as per-sample coverage increases. Therefore, there may be higher levels of coverage for which Boiler shuffling has a greater impact than technical-replicate shuffling. For the realistic levels of coverage we tested, however, Boiler’s shuffling remains less consequential.

We also measured precision and recall in an unweighted fashion, i.e. omitting the coverage terms *c(t)* and 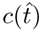 from equations 2 and 3. These results are similar to the weighted results as discussed in Supplementary Note 8.

### Queries

Recovering coverage vectors from a Boiler-compressed file requires that Boiler decompress and combine coverage vectors for all the relevant bundles and buckets. Decompression of the coverage vector involves the DEFLATE algorithm and run-length decoding, but does not involve the more expensive read and pair recovery algorithms. SAMtools, on the other hand, does not explicitly represent the coverage vectors in a BAM file. Instead, coverage information must be recovered from the BAM file by first extracting the relevant alignments, then composing the coverage vector using another tool like BEDTools: 

~~~
samtools view -b -h x.bam c:start-end | genomeCoverageBed -bga -split -ibam stdin -g chromosomes.txt
~~~

**Table 7.** Reference-based Recall

| Dataset | Cufflinks |  | StringTie |  | HISAT + StringTie |  |
| --- | --- | --- | --- | --- | --- | --- |
|  | Original | Compressed | Original | Compressed | Original | Compressed |
| Drosophila, Simulated Unpaired |  |  |  |  |  |  |
| 0.5M | 0.583 | 0.583 (+0.0%) | 0.526 | 0.526 (+0.0%) | 0.551 | 0.551 (+0.0%) |
| 1M | 0.708 | 0.707 (-0.1%) | 0.682 | 0.682 (+0.0%) | 0.712 | 0.712 (+0.0%) |
| 2.5M | 0.791 | 0.785 (-0.7%) | 0.795 | 0.795 (-0.0%) | 0.803 | 0.800 (-0.4%) |
| 5M | 0.822 | 0.822 (+0.0%) | 0.834 | 0.834 (-0.0%) | 0.833 | 0.831 (-0.3%) |
| 10M | 0.824 | 0.824 (+0.0%) | 0.844 | 0.844 (+0.0%) | 0.835 | 0.834 (-0.2%) |
| 20M | 0.827 | 0.826 (-0.1%) | 0.851 | 0.850 (-0.1%) | 0.853 | 0.847 (-0.7%) |
| Drosophila, Simulated Paired |  |  |  |  |  |  |
| 0.5M | 0.732 | 0.729 (-0.4%) | 0.691 | 0.689 (-0.3%) | 0.715 | 0.715 (-0.1%) |
| 1M | 0.799 | 0.797 (-0.2%) | 0.792 | 0.791 (-0.1%) | 0.814 | 0.814 (-0.0%) |
| 2.5M | 0.825 | 0.826 (+0.1%) | 0.840 | 0.840 (-0.0%) | 0.843 | 0.840 (-0.4%) |
| 5M | 0.840 | 0.838 (-0.2%) | 0.857 | 0.855 (-0.2%) | 0.861 | 0.859 (-0.3%) |
| 10M | 0.828 | 0.824 (-0.5%) | 0.851 | 0.851 (-0.0%) | 0.851 | 0.849 (-0.3%) |
| 20M | 0.826 | 0.823 (-0.4%) | 0.850 | 0.849 (-0.1%) | 0.854 | 0.849 (-0.6%) |
| Human, Simulated Paired |  |  |  |  |  |  |
| 20M | 0.762 | 0.762 (+0.0%) | 0.794 | 0.792 (-0.2%) | 0.851 | 0.850 (-0.1%) |
| 40M | 0.792 | 0.789 (-0.3%) | 0.824 | 0.821 (-0.4%) | 0.851 | 0.849 (-0.2%) |

We compared the time required for Boiler to respond to coverage queries to the time required for SAMtools/BEDTools. Specifically, we iterated over all bundle boundaries in several *D. melanogaster* samples and queried for the coverage vector within those boundaries using both Boiler and SAMtools/BEDTools. Figure 5 compares the tools both in terms of average query time (left) and per-bundle query time (right). As expected, Boiler is consistently faster than SAMtools/BEDTools, with Boiler taking under 0.1 seconds on average, and SAMtools/BEDTools taking close to 0.75 seconds.

**Table 8.** Non-reference-based precision. Columns labeled Boiler compare precision before and after Boiler compression. Columns labeled Tech Reps compare pairs of technical replicates.

| Dataset | Cufflinks |  | Stringtie |  |
| --- | --- | --- | --- | --- |
|  | Boiler | Tech Reps<br>(min–max) | Boiler | Tech Reps<br>(min–max) |
| Drosophila, Simulated Unpaired |  |  |  |  |
| 0.5M | 0.997 | 0.386–0.396 | 0.998 | 0.452–0.470 |
| 1M | 0.996 | 0.492–0.502 | 0.998 | 0.559–0.578 |
| 2.5M | 0.989 | 0.659–0.669 | 0.993 | 0.729–0.736 |
| 5M | 0.988 | 0.757–0.763 | 0.989 | 0.807–0.816 |
| 10M | 0.983 | 0.814–0.824 | 0.985 | 0.854–0.863 |
| 20M | 0.980 | 0.861–0.868 | 0.983 | 0.892–0.900 |
| Drosophila, Simulated Paired |  |  |  |  |
| 0.5M | 0.986 | 0.618–0.636 | 1.000 | 0.568–0.583 |
| 1M | 0.980 | 0.724–0.733 | 0.996 | 0.698–0.706 |
| 2.5M | 0.965 | 0.806–0.811 | 0.989 | 0.807–0.815 |
| 5M | 0.969 | 0.843–0.850 | 0.991 | 0.854–0.862 |
| 10M | 0.960 | 0.867–0.879 | 0.986 | 0.890–0.895 |
| 20M | 0.952 | 0.893–0.902 | 0.985 | 0.915–0.921 |
| Human |  |  |  |  |
| SRP025982 (11M) | 0.964 | 0.627–0.645 | 0.997 | 0.622–0.641 |
| HG00100 (20M) | 0.939 | (no replicates) | 0.997 | (no replicates) |
| Simulated 20M | 0.969 | 0.897–0.912 | 0.981 | 0.910–0.932 |
| Simulated 40M | 0.965 | 0.911–0.927 | 0.976 | 0.935–0.947 |

**Figure 5.**
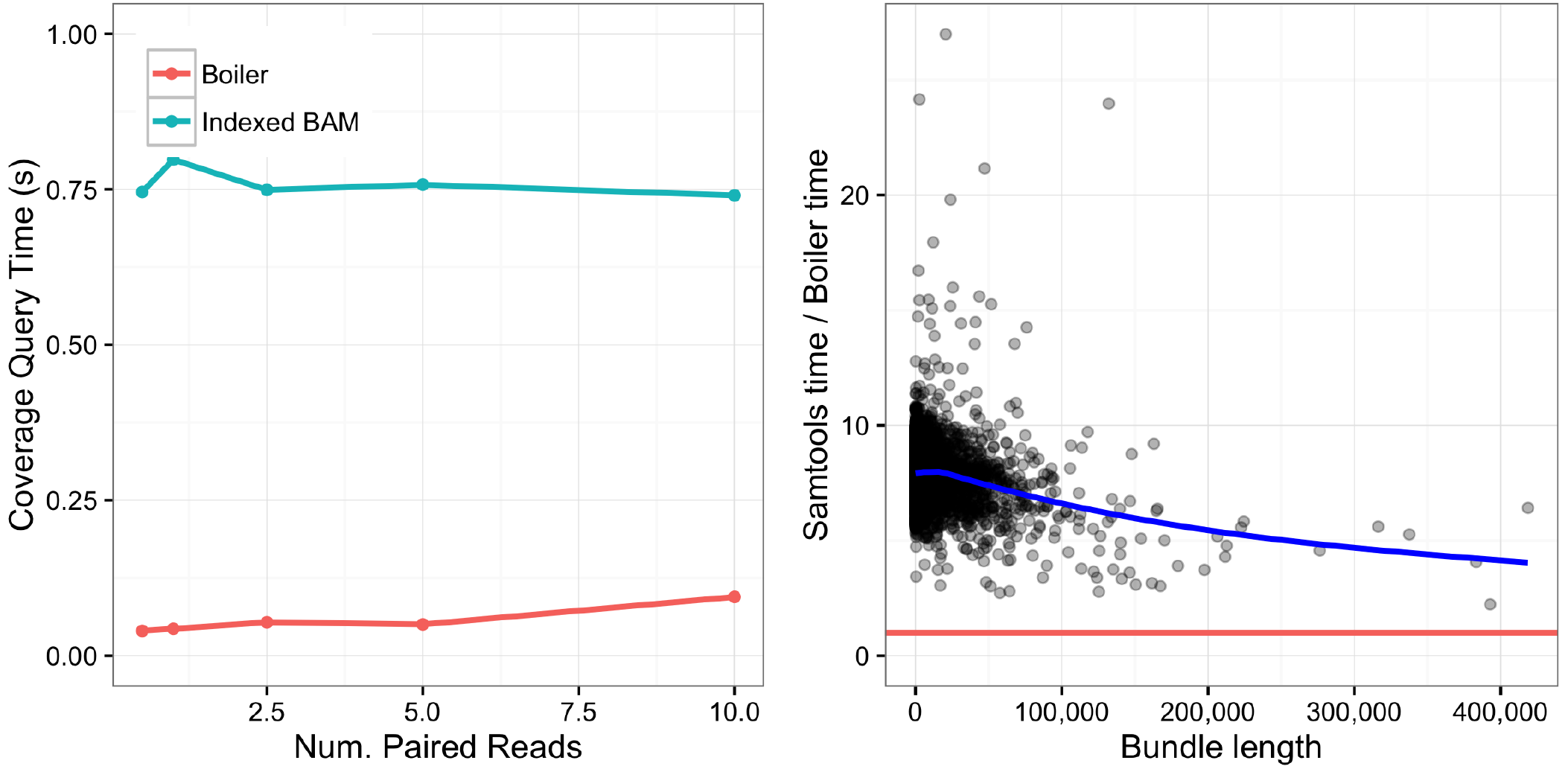
Comparison of coverage query times for Boiler the indexed BAM for all bundles. Left: Average query time for varying *D. melanogaster* paired-end datasets. Right: Ratio of Samtools / Boiler query time for each bundle in the 10M *D. melanogaster* paired-end dataset plotted as a function of bundle length. The blue line denotes the best fit line for the points, the red line is *y* = 1.

We also compared alignment query times for Boiler to those for the indexed BAM. A SAMtools alignment query uses this command: 

~~~
samtools view -h x.bam chrom:start-end
~~~

Boiler must both decompress the relevant bundles and run the greedy read and pair recovery algorithms. Figure 6 compares the tools both in terms of average query time (left) and per-bundle query time (right). As expected, because of the need to run read and pair recovery, Boiler’s alignment query is consistently slower than SAMtools. Even so, Boiler’s average response time is under 0.15 seconds.

**Figure 6.**
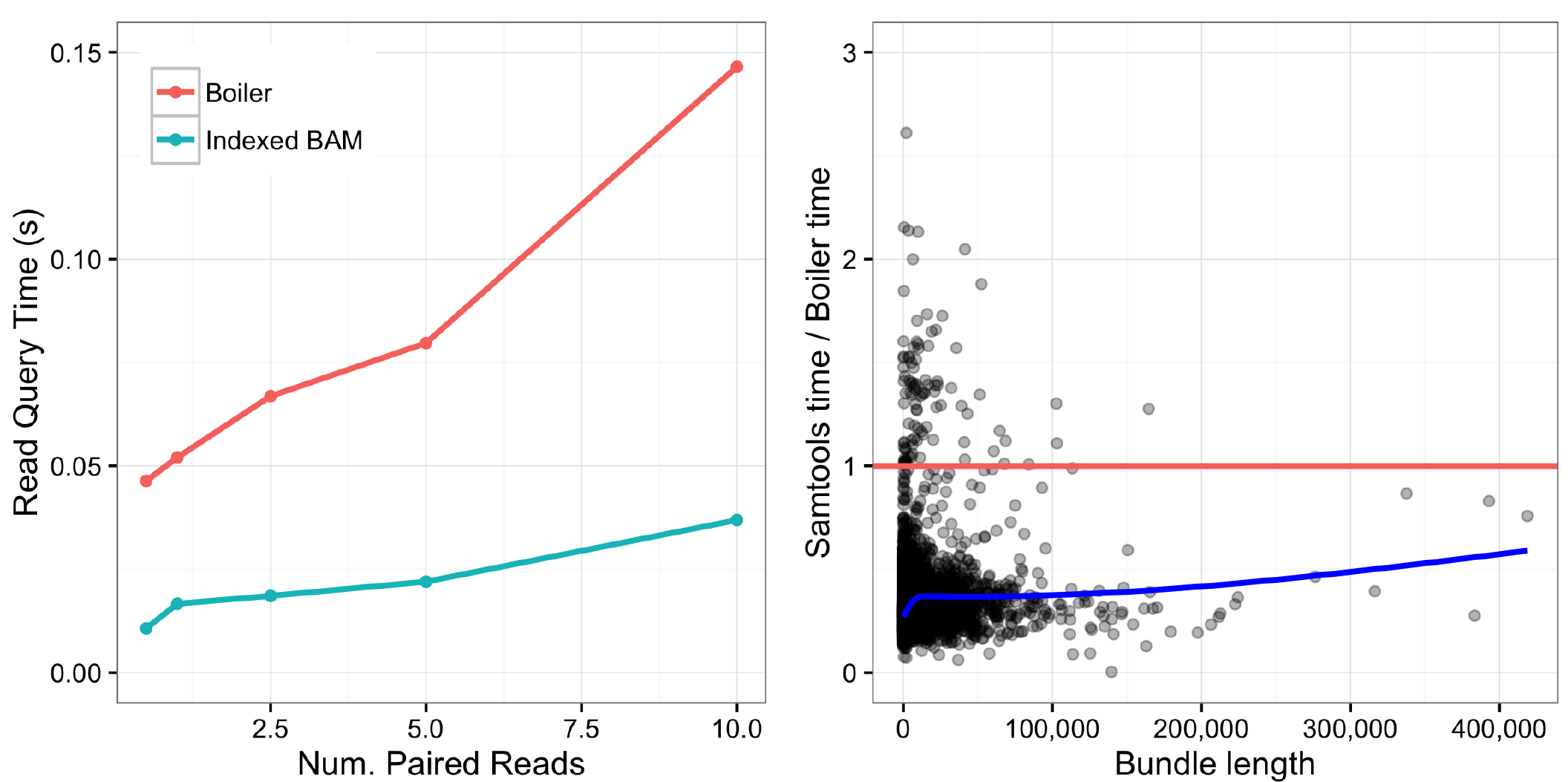
Comparison of alignment query times for Boiler versus sorted and indexed BAM for all bundles. Left: Average query time for varying *D. melanogaster* paired-end datasets. Right: Ratio of Samtools / Boiler query time for each bundle in the 10M *D. melanogaster* paired-end dataset plotted as a function of bundle length. The blue line denotes the best fit line for the points, the red line is *y* =1.

### DISCUSSION

Boiler applies principles of lossy compression and transform coding to the problem of compressing RNA-seq alignments. Beyond discarding unnecessary BAM attributes, Boiler additionally discards most of the data that ties individual reads to their aligned positions and shapes. Boiler instead stores coverage vectors and read- and outer-distance tallies, effective shifting from the “alignment domain” to the “coverage domain.” While this can cause alignments to shift along the genome or pair with the wrong mate, the shuffling effect is modest compared to the shuffling induced by switching between technical replicates, and adverse effects on downstream tools for isoform assembly and quantification are minimal.

**Table 9.**
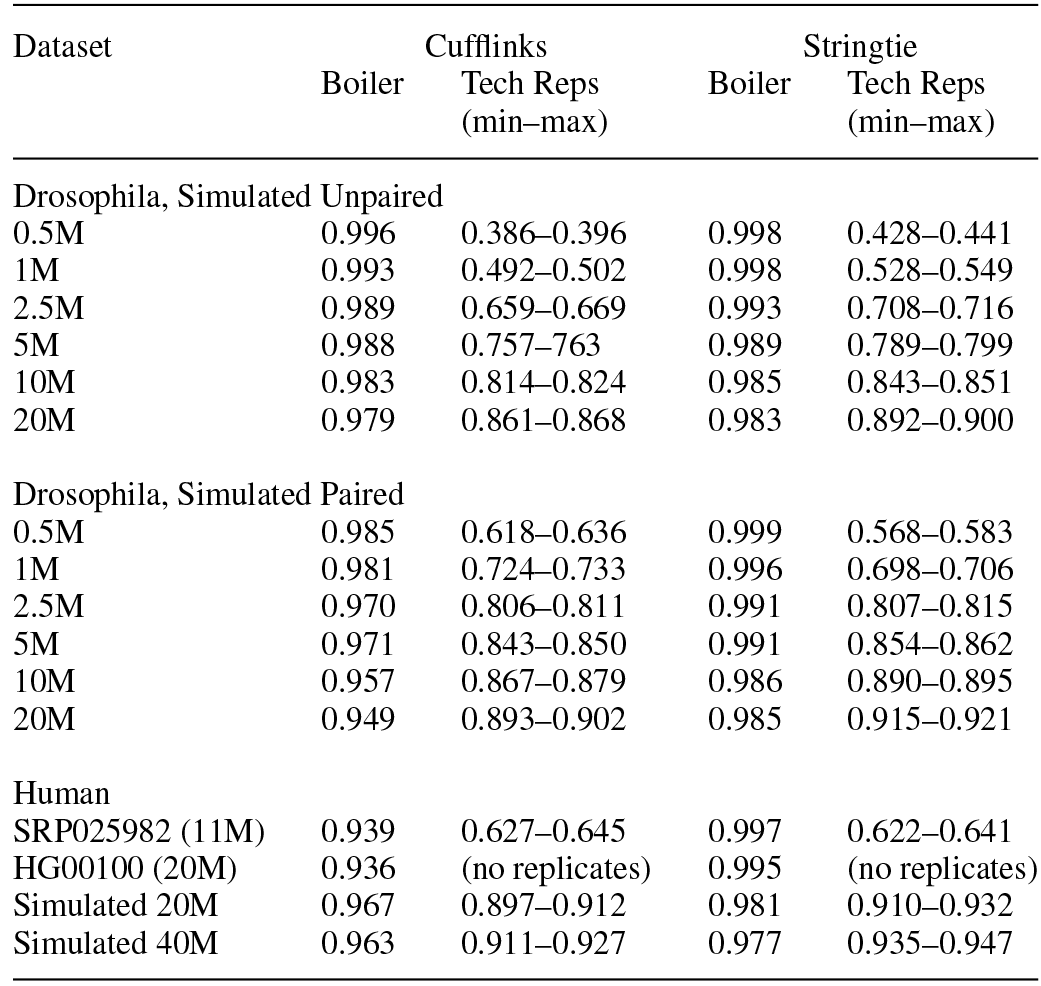
Non-reference-based recall. Columns labeled Boiler compare recall before and after Boiler compression. Columns labeled Tech Reps compare pairs of technical replicates.

| Dataset | Cufflinks |  | Stringtie |  |
| --- | --- | --- | --- | --- |
|  | Boiler | Tech Reps<br>(min–max) | Boiler | Tech Reps<br>(min–max) |
| Drosophila, Simulated Unpaired |  |  |  |  |
| 0.5M | 0.996 | 0.386–0.396 | 0.998 | 0.428–0.441 |
| 1M | 0.993 | 0.492–0.502 | 0.998 | 0.528–0.549 |
| 2.5M | 0.989 | 0.659–0.669 | 0.993 | 0.708–0.716 |
| 5M | 0.988 | 0.757–0.763 | 0.989 | 0.789–0.799 |
| 10M | 0.983 | 0.814–0.824 | 0.985 | 0.843–0.851 |
| 20M | 0.979 | 0.861–0.868 | 0.983 | 0.892–0.900 |
| Drosophila, Simulated Paired |  |  |  |  |
| 0.5M | 0.985 | 0.618–0.636 | 0.999 | 0.568–0.583 |
| 1M | 0.981 | 0.724–0.733 | 0.996 | 0.698–0.706 |
| 2.5M | 0.970 | 0.806–0.811 | 0.991 | 0.807–0.815 |
| 5M | 0.971 | 0.843–0.850 | 0.991 | 0.854–0.862 |
| 10M | 0.957 | 0.867–0.879 | 0.986 | 0.890–0.895 |
| 20M | 0.949 | 0.893–0.902 | 0.985 | 0.915–0.921 |
| Human |  |  |  |  |
| SRP025982 (11M) | 0.939 | 0.627–0.645 | 0.997 | 0.622–0.641 |
| HG00100 (20M) | 0.936 | (no replicates) | 0.995 | (no replicates) |
| Simulated 20M | 0.967 | 0.897–0.912 | 0.981 | 0.910–0.932 |
| Simulated 40M | 0.963 | 0.911–0.927 | 0.977 | 0.935–0.947 |

Boiler is not a general-purpose substitute for RNA-seq SAM/BAM files, but it is an extremely space-efficient alternative that works well with tools like Cufflinks and StringTie. RNA-seq alignments are much larger than downstream files summarizing coverage (BigWig) or per-isoform expression level (FPKM table). Boiler and BigWig are similar in size, but, importantly, Boiler files preserve the ability to re-run the analysis repeatedly in the future. This can profoundly reduce the cost and difficulty of working with RNA-seq data, especially for large datasets.

Though we have not explored it here, we expect Boiler compression to work well with annotation-based tools like featureCounts (18) and HiTseq (1), as well as downstream differential-expression tools like derfinder (8) that rely on counts and coverage values.

Boiler also discards information about non-reference alleles, making Boiler archives readily sharable even when the input data is protected by privacy provisions like dbGaP.

In future work, it will be important to explore alternatives to the greedy decompression algorithms described here. For example, Boiler’s greedy algorithm for extracting reads from a coverage distribution assumes the distribution of input read lengths has a dominant mode. This is a reasonable assumption for Illumina data, but not so for other sequencing technologies. Also, Boiler’s compression and decompression algorithms could be accelerated by moving from Python to a compiled language such as C/C++. In general, there are many opportunities to make the read extraction and pairing algorithms more accurate, faster, and less hampered by assumptions about the data.

Another subject for future work is how Boiler represents multi-mapping alignments. By discarding read names, Boiler discards the one-to-many relationship between a multimapping read and its alignments. While this does not harm fidelity in most cases, it does adversely affect fidelity when Cufflinks is used to quantify from a gene annotation. It is an open question as to whether and how multi-mapping relationships can be represented in a way that allows trading off between compression ratio and fidelity.

Finally, we note that the methods used here to characterize Boiler’s shuffling effect are more generally useful for evaluating any upstream tool that modifies the data. For example, one could apply the same techniques to evaluate a tool for read trimming or digital normalization, or to compare many parameterizations of the spliced aligner.

Boiler is available from github.com/jpritt/boiler and is distributed under the open source MIT license.

## ACKNOWLEDGEMENTS

We are grateful to Abhinav Nellore and Chris Wilks for their comments on the software and manuscript. We are also grateful to Fabien Campagne for his comments on the manuscript.

### FUNDING

JP was supported and BL was partially supported by a Sloan Research Fellowship to BL. BL was partially supported by National Science Foundation (IIS-1349906) and National Institute of General Medical Sciences (1R01GM118568) grants to BL.

## REFERENCES

1. AndersS, PylPT, HuberW (2014) HTSeq-A Python framework to work with high-throughput sequencing data. Bioinformatics, 31(2), 166–169.

2. ArdlieKG, DelucaDS, SegrèAV, SullivanTJ, YoungTR, GelfandET, TrowbridgeCA, MallerJB, TukiainenT, LekM et al. (2015) The Genotype-Tissue Expression (GTEx) pilot analysis: Multitissue gene regulation in humans. Science, 348(6235), 648–660.

3. BonfieldJK, MahoneyMV (2013) Compression of fastq and sam format sequencing data. PLoS ONE, 8(3), e59190.

4. CampagneF, DorffK, ChambweN, RobinsonJT, MesirovJP (2013) Compression of Structured High-Throughput Sequencing Data. PLoS ONE, 8(11), e79871.

5. Cufflinks pull request 32, https://github.com/cole-trapnell-lab/cufflinks/pull/32 (2015).

6. DailyK, RigorP, ChristleyS, XieX, BaldiP (2010) Data structures and compression algorithms for high-throughput sequencing technologies. BMC bioinformatics, 11(1), 514.

7. FilippovaD, KingsfordC (2015) Rapid, separable compression enables fast analyses of sequence alignments. ACM Conference on Bioinformatics, 194–201.

8. FrazeeA, SabunciyanS, HansenK, IrizarryR, LeekJ (2014) Differential expression analysis of RNA-seq data at single-base resolution. Biostatistics, 15(3), 413–426.

9. HachF, NumanagicI, AlkanC, SahinalpSC (2012) SCALCE: boosting sequence compression algorithms using locally consistent encoding. Bioinformatics, 28(23), 3051–3057.

10. HarrowJ, FrankishA, GonzalezJM, TapanariE, DiekhansM, KokocinskiF, AkenBL, BarrellD, ZadissaA, SearleS et al. (2012) GENCODE: the reference human genome annotation for The ENCODE Project. Genome research, 22(9), 1760–1774.

11. Hsi-YangFM, LeinonenR, CochraneG, BirneyE (2011) Efficient storage of high throughput DNA sequencing data using reference-based compression. Genome Research, 5, 734–740.

12. JonesDC, RuzzoWL, PengX, KatzeMG (2012) Compression of next-generation sequencing reads aided by highly efficient de novo assembly. Nucleic Acids Research, 40(22), e171.

13. KentWJ, ZweigAS, BarberG, HinrichsAS, KarolchikD (2010) BigWig and BigBed: enabling browsing of large distributed datasets. Bioinformatics, 26(17), 2204–2207.

14. KimD, PerteaG, TrapnellC, PimentelH, KelleyR, SalzbergSL (2013). TopHat2: accurate alignment of transcriptomes in the presence of insertions, deletions and gene fusions Genome Biol, 14(4), R36.

15. KimD, LangmeadB, SalzbergSL (2013). HISAT: a fast spliced aligner with low memory requirements Nature methods, 12(4), 357–360.

16. KozanitisC, SaundersC, KruglyakS, BafnaV, VargheseG (2011) Compressing genomic sequence fragments using SlimGene. Journal of Computational Biology, 18(3), 401–413.

17. LappalainenT, SammethM, FriedländerMR, AC?t HoenP, MonlongJ, RivasMA, Gonzàlez-PortaM, KurbatovaN, GriebelT, FerreiraPG et al. (2013) Transcriptome and genome sequencing uncovers functional variation in humans. Nature, 501(7468), 506–511.

18. LiaoY, SmythGK, ShiW (2014) featureCounts: an efficient general purpose program for assigning sequence reads to genomic features. Bioinformatics, 30(7), 923–930.

19. GriebelT, ZacherB, RibecaP, RaineriE, LacroixV, GuigoR, SammethM (2012) Modelling and simulating generic RNA-Seq experiments with the flux simulator. Nucleic acids research, 40(20), 10073–10083.

20. LeinonenR, SugawaraH, ShumwayM (2011) The sequence read archive. Nucleic acids research, 39(suppl 1), D19–D21.

21. LiH, DurbinR (2009) Fast and accurate short read alignment with Burrows-Wheeler Transform. Bioinformatics, 25, 1754–1760.

22. LiH, HandsakerB, WysokerA, FennellT, RuanJ, HomerN, MarthG, AbecasesG, DurbinR (2009) The Sequence Alignment/Map format and SAMtools. Bioinformatics, 25(16), 2078–2079.

23. OchoaI, AsnaniH, BharadiaD, ChowdhuryM, WeissmanT, YonaG (2013) QualComp: a new lossy compressor for quality scores based on rate distortion theory. BMC Bioinformatics, 14, 187.

24. PerteaM, PerteaGM, AntonescuCM, ChangTS, MendellJT, SalzbergSL (2015). StringTie enables improved reconstruction of a transcriptome from RNA-seq reads. Nature Biotechnology, 33(3), 290–295.

25. PopitschN, vonHaeseler A (2013). NGC: lossless and lossy compression of aligned high-throughput sequencing data. Nucleic Acids Research, 41(1), e27.

26. SEQC/MAQC-III Consortium (2014) A comprehensive assessment of RNA-seq accuracy, reproducibility and information content by the Sequencing Quality Control Consortium. Nat Biotechnol, 32(9), 903–914.

27. StephensZD, LeeSY, FaghriF, CampbellRH, ZhaiC, EfronMJ, IyerR, SchatzMC, SinhaS, RobinsonGE (2015) Big Data: Astronomical or Genomical? PLoS Biol, 13(7), e1002195.

28. TrapnellC, WilliamsB, PerteaG, MortazaviA, KwanG, vanBaren J, SalzbergS, WoldB, PachterL (2010).Transcript assembly and quantification by RNA-Seq reveals unannotated transcripts and isoform switching during cell differentiation. Nature Biotechnology, 28(5), 511–515.

29. YatesA, AkanniW, AmodeMR, BarrellD, BillisK, Carvalho-SilvaD, CumminsC, ClaphamP, FitzgeraldS, GilL et al. (2015). Ensembl 2016 Nucleic Acids Research, 28(5), gkv1157.

